# A deep learning algorithm for potato tuber hollow heart classification

**DOI:** 10.1101/2021.09.29.462243

**Authors:** Arash Abbasi, Max J. Feldman, Jaebum Park, Katelyn Greene, Richard G. Novy, Huaping Liu

## Abstract

A novel deep learning algorithm is proposed for hollow heart detection which is an internal tuber defect. Hollow heart is one of many internal defects that decrease the market value of potatoes in the fresh market and food processing sectors. Susceptibility to internal defects like the hollow heart is influenced by genetic and environmental factors so elimination of defect-prone material in potato breeding programs is important. Current methods of evaluation utilize human scoring which is limiting (only collects binary data) relative to the data collection capacity afforded by computer vision or are based upon X-ray transmission techniques that are both expensive and can be hazardous. Automation of defect classification (e.g. hollow heart) from data sets collected using inexpensive, consumer-grade hardware has the potential to increase throughput and reduce bias in public breeding programs. The proposed algorithm consists of ResNet50 as the backbone of the model followed by a shallow fully connected network (FCN). A simple augmentation technique is performed to increase the number of images in the data set. The performance of the proposed algorithm is validated by investigating metrics such as precision and the area under the curve (AUC).

## I. Introduction

Potato (*Solanum tuberosum*) is the most economically important vegetable crop in the United States. Over the last decade, the U.S. has consistently planted over 900, 000 acres of potato, resulting in the production of over 400 million CWT of tuber material and an economic value approaching four billion U.S. dollars [1]. Roughly two-thirds of the crop is sold to the food processing industry to generate products like French fries, potato chips, dehydrated potato products, and canned goods, whereas a quarter is sold as a fresh market product to consumers in grocery stores and restaurants. Quality standards for potatoes are rigorous in both the processing and fresh market industry. Defects can occur during the growth season (hollow heart, internal browning syndrome, growth cracks, virus-induced necrotic lesions), harvest (bruise, cracking, greening), and storage (sprouting, greening); the consequence of which is a financial loss to the grower. Reducing the propensity of cultivars to express these defects is one major objective of potato breeding programs. In some cases, genetic methods can be used to predict and select against offspring that will be susceptible to defects. Unfortunately, tabulating the type and severity of tuber defects on the scale needed to evaluate early-stage breeding populations (> 200 clones; between 5 to 100 tubers per clone) is not trivial. Challenges arise as symptomology can be similar between defect type and time which can be allocated to observe any individual tuber is very small. The low-cost, high information content, and rapid acquisition time make digital imaging of tubers an efficient way to document the properties off-spring in breeding populations.

With the rapid increase in computation power and big data, deep learning has shown tremendous success in various image and signal processing applications such as health care, biology, anomaly detection [2, 3, 4, 5, 6]. Unlike conventional rule-based algorithms, in deep learning algorithms suitable discriminatory features for classification/detection are extracted by the model automatically during the training phase. Hence, better performance is achievable.

In this report, a novel deep learning approach is proposed for potato hollow heart detection. Hollow heart is a physiological, internal tuber defect that can occur when growth in the perimedullary region of the tuber outpaces growth of the pith causing the development of lens-, star-, or irregularly-shaped cavities within the pith tissue [7]. Frequently, expression of hollow heart is a consequence of water or nutrient fluctuation during the growth season [7]. Genetics has been demonstrated to play a role in hollow heart susceptibility [8] and in some cases susceptibility is likely to be independent of final tuber size [9].

The presence of the hollow heart is somewhat common and relatively easy to identify, making it an excellent test case to evaluate the feasibility of applying deep learning approaches to defect classification in plant breeding programs. A hollow heart and non-hollow heart examples are shown in Fig. 1 and Fig. 2, respectively.

**Fig. 1:**
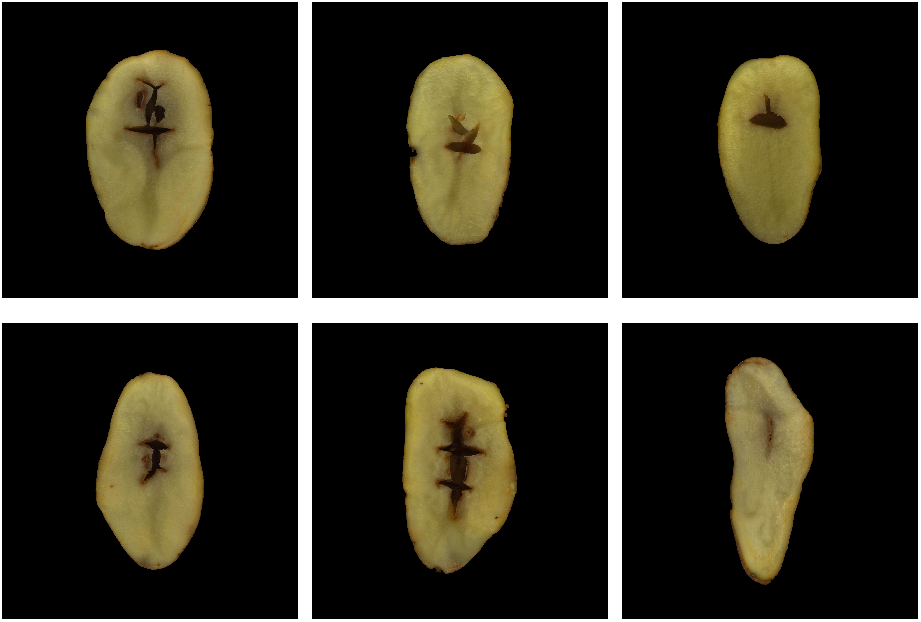
Hollow heart.

**Fig. 2:**
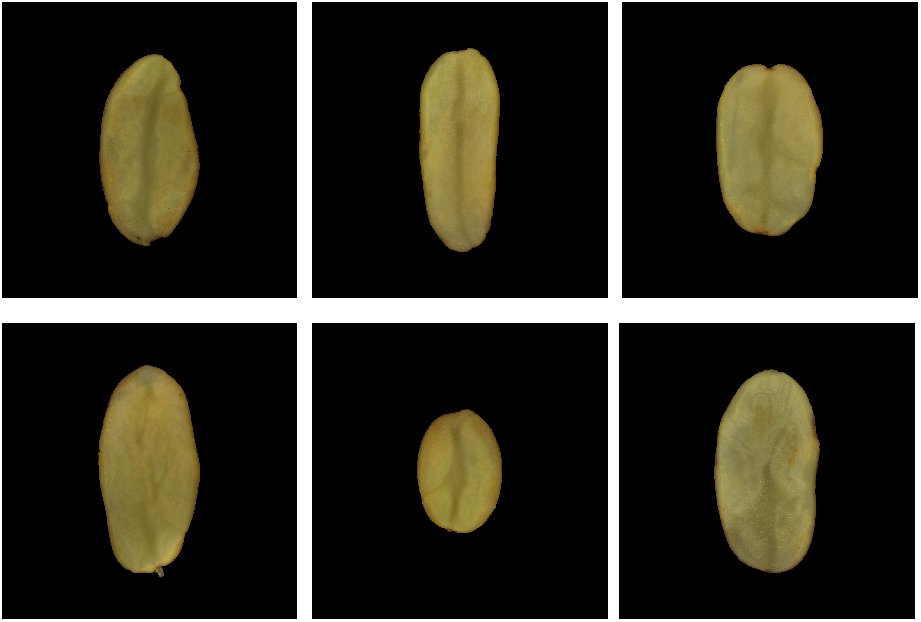
Non-hollow heart.

## II. Methods

The data set contains 1610 three-channel RGB images of potato tuber cut in half lengthwise. Each image is 1900 × 1900 pixels x pixels, contains a single tuber that is centered on a black background. The presence of internal tuber defects such as hollow heart, internal brown spot, bruise, anthocyanin accumulation, and tuber greening was scored manually according to a USDA visual aid guide provided in [10].

### A. Plant material

The subjects are tubers derived from 189 F1 progeny from the A08241 autotetraploid linkage mapping population. In 2019, both parental lines (Palisade Russet, female; ND02873 2Russ, male) and their progeny were planted in a randomized complete block design with two replications of eight-hill plots by scientists at the USDA-ARS Small Grains and Potato Germplasm Research Unit in Aberdeen, ID.

### B. Digital imaging of tubers

Computer vision measurements are acquired by scientists at the USDA-ARS Temperate Tree Fruit and Vegetable Research Unit in Prosser, WA. Digital scans of tuber internal characteristics are acquired using a Hewlett Packard HP ColorJet 6200C flatbed scanner (See Supplemental Fig. 1). Each clone was replicated twice, and each replicate contains five tubers within the image. Before scanning each tuber is cut in half lengthwise and an image of each tuber is captured. Both size and radiometric calibration standards are included in each image. A blue plastic poker chip (37 mm diameter) is used as a size standard in all images and scans. An X-Rite ColorChecker Classic is used as a radiometric standard for the top-down imaging on the black background whereas an X-Rite ColorChecker Mini is used as color standards for the scanner. Python scripts used to control the imaging hardware can be found at [11]. Individual tubers were labeled by their position in the image and manually scored for the presence or absence of internal defects according to the USDA grading standards visual aid chart.

## III. Proposed Deep Learning Architecture

### A. Optimization

The goal of the proposed algorithm is to identify hollow heart defects in the potato images. This can be sought as a binary classification task, i.e., hollow heart vs non-hollow heart classification, and hence deep learning algorithms suitable for classification applications can be employed. Let’s *C*_0_ and *C*_1_ represent the non-hollow heart and hollow heart classes, respectively, and d an image in the data set. In this case, we aim to find the probability that the image *d* is a member of non-hollow heart *P(C*_0_ |*d*) or hollow-heart *P(C*_1_ |*d*) sets. Since there are two discrete outcome, the Bernoulli distribution function can be derived to express the conditional probability *P*(*y |x*), that is the probability of the the image class is y for an observation x, as

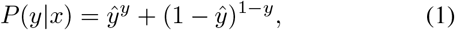

where *y* ϵ {0, 1} is the true label and *ŷ* is the model prediction for the image *x*. The log likelihood estimation can be employed that aims to maximize the log *p*(*y* |*x*). In this case, the log-likelihood cost-function C_*LL*_ is defined as

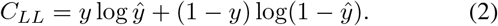

It is seen that maximizing the log-likelihood cost-function,i.e., maximizing the probability of the correct estimated labels, is equivalent to minimizing the cross-entropy between true labels and the estimated labels which is defined as

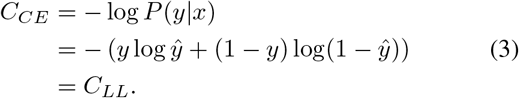

This ensures the maximization of the correct prediction and minimization of the incorrect classification. Assuming there are *M* independent and identically distributed (iid) images in the training data set, the log-likelihood cost-function for the training data set is given as

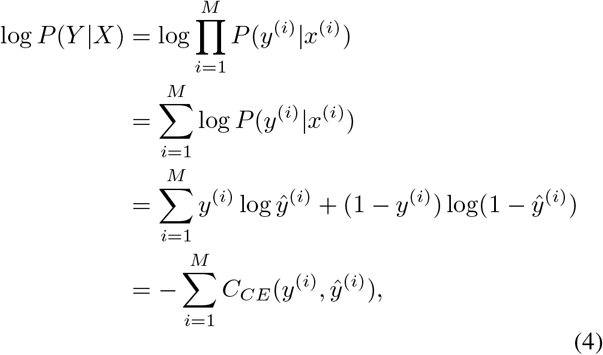

where *Y* = {*y*_1_, *y*_2_, …, *y*_*M*_} and *X* = {*x*_1_, *x*_2_, …, *x*_*M*_} are the sets of all labels and images in the training data sets. By employing deep learning algorithms, the cross-entropy cost-function in (4) is parametrized by the weights of the model Θ = (*θ*^(1)^, *θ*^(2)^,…, *θ*^(*L*)^), in which L represents the number of hidden layers in the model. We represent the output of the model as *f*(*x*^(*i*)^; Θ) for the *i*-th image *x*^(*i*)^. In this case, the optimum solution for the weight of the model Θ, weights that minimized the cross-entropy cost-function, can be given as

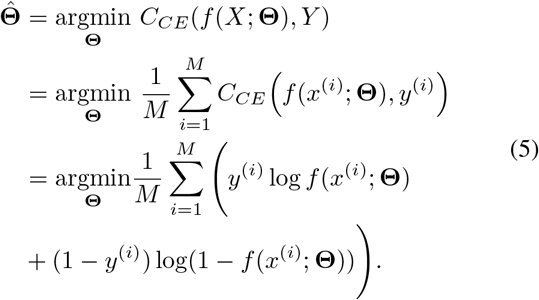

To minimize potential overfitting and address the sparsity in the data set, *𝓁*_2_ and *𝓁*_1_ regularization are employed, respectively. In this case, the cost-function can be written as

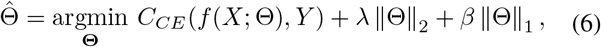

where λ and α are regularization coefficients for *𝓁*_2_ and *𝓁*_1_ regularization, respectively. Various gradient descent algorithms can be employed that aim to find the set of In this paper, adaptive moment estimation (Adam) is used to find the set of the optimum solution for the weights of the model Θ. in (6), in terms of minimizing cross-entropy or maximizing the log-likelihood [13].

### B. Proposed Deep Learning Model

In this paper, we propose two deep learning architectures that are composed of ResNet50 [12] followed by different shallow fully connected networks (FCNs). A block diagram of the proposed architectures is depicted in Figure 3. The shallow FCN is added to the ResNet50 to decrease the dimensionality of the last layer of the ResNet50 so that it will be suitable for binary classification. The proposed architectures are summarized in Table I. The cross-entropy (CE) loss derived in (4, the area under the curve, and the precision of the models are evaluated. Precision of the proposed modes is measured as

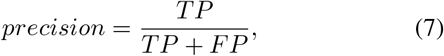

where TP and FP are true positive and false positive respectively. The categorical accuracy of the model is calcualted as

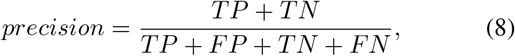

where TN and FN are true negative and false negative, respectively.

**Fig. 3:**
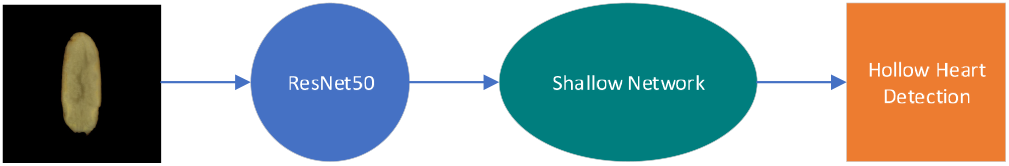
A simple block diagram of the proposed model.

**TABLE I:** The proposed model architectures.

|  |  |
| --- | --- |
| Model 1 | ResNET50 → (Dense(256), ReLU) → (Dense(128), ReLU) $\times 3 \rightarrow$ (Dense(128), sigmoid) → (Dense(2), sigmoid) |
| Model 2 | ResNET50 → (Dense(128), sigmoid) → (Dense(2), sigmoid) |

## IV. Simulation Results

The data set consists of 50 potato images with the hollow heart defect and 1730 images with no hollow heart. It is obvious that the data set is unbalanced. Therefore in a prepossessing step, we aim to 1) balance the data set by selecting randomly only 50 images from the non-hollow heart defect class and 2) increase the number of the images in the data set by performing a simple augmentation procedure that is rotating the images by 90 and 180 degrees and adding additive white Gaussian noise (AWGN) with *µ* =0 and variance σ^2^. After the prepossessing step, there are 150 potato images with hollow-heart deficiency and 150 with no hollow heart deficiency in the training data set. Therefore, in total there are 300 images in the data set.

There are 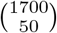 different permutations for selecting 50 images from total 1700 non-hollow heart images. To increase the model exposure to more diverse non-hollow cases, a Monte Carlo approach is employed where the model is trained 120 times independently. The dimension of the potato images is 1200 × 1200 pixels x pixels. The images are resized to 100 × 100 pixels x pixels to reduce the redundancy in the images and enhance the computational processing speed.

Additionally, since the subject is plotted on a completely black background in each image and the background will not influence defect status, a cropping mechanism is employed so that only the potato portion of the images is considered in the cropped images and the background is removed from each image

The cross-entropy loss, the precision, the area under the curve (AUC), and the categorical accuracy performance of the proposed algorithms when the variance of the AWGN is σ^2^ = {0.01, 0.03, 0.05} and for different regularization, values are depicted in Figs. 4. The learning rate, regularization parameter, and the batch size are {1e−3, 1e−5, 50}, respectively. As expected, the performance of the model in the training phase outperforms the validation performance. Moreover, it is seen that the model 1 outperforms the model 2. This demonstrates the effectiveness of the deeper FCN added to the ResNet model. In other words, as the depth of the FCN increases, the model can learn more discrepancy features from the input images and hence the performance of the model improves. Additionally, for all three metrics, the performance of the models improves as the variance of the AWGN decreases. This is because, in lower variance, the model can learn better discrepancy features from the images, and hence, better performance can be achieved. Surprisingly, however, it is seen that the performance of the models degrade when regularization is added to the models. One hypothesis for this anomaly behavior is that since the number of images in the training data set is small, hence, the added regularization terms behave restrictive for the model weights that are set to be trained. It is also seen that the categorical accuracy performance is identical to the precision. This is because the model is balanced. That is, its ability to correctly classify hollow heart (positive) samples is same as its ability to correctly classify non-hollow heart (negative) samples. The accuracy and precision performance demonstrate the capacity of the model to detect and identify the hollow heart (and therefore, non-hollow heart) cases. For example, after 800 training epochs, the model 1 can achieve 90% accuracy.

**Fig. 4:**
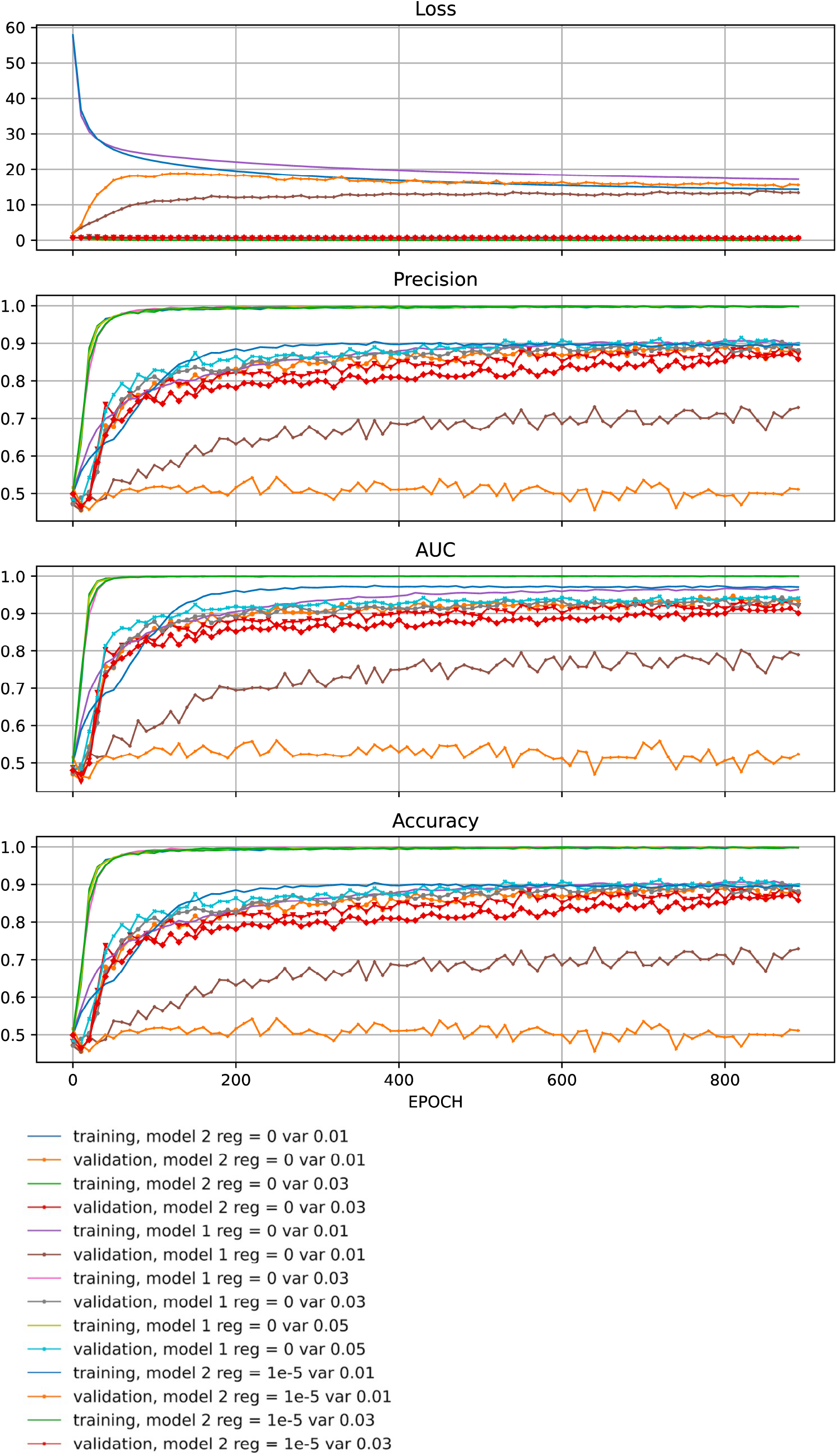
The performance of the proposed models.

## V. Discussion

This study suggests that deep learning techniques applied to image data sets collected using consumer grade hardware may be sufficient to classify the presence or absence of tuber defects in breeding populations; so long as the training set is representative of the classification problem and contains enough images to train the model. As demonstrated, augmentation of data classes using image rotation and introduction of additive white Gaussian noise may be used to increase the size of the training set but caution is advised in application settings. The model described may not perform as expected if applied to other data sets, particularly data sets containing tubers with different shapes and/or flesh colors.

Despite the need for destructive sampling, the hollow heart classification approach presented in this study offers several advantages over other commonly used diagnostic methods described in the literature. The relationship between tuber size and specific gravity has been used identify and remove potentially affected tubers in bulk grading applications [14, 15, 16]. Unfortunately, manual destructive assessment is still required to achieve the precision needed for breeding applications. Although relatively high precision has been achieved using light transmittance [17], confounding variation associated with differences in tuber diameter, tuber skin thickness, and tuber skin quality make the evaluation of the diverse tuber material encountered in breeding populations challenging. The application of non-destructive ultrasonic technologies is fraught with difficulties due to the high attenuation coefficient of potato tubers, need for multiple transducers, and frictional noise generated by the tuber moving along the transducer [21]. Excellent success of hollow heart detection and sample throughput has been achieved using non-destructive techniques including X-ray machines [14, 15, 18, 19] and hyperspectral sensors [20]. Unfortunately, the equipment (X-ray and hyperspectral) and safety measures (X-ray) required for deployment make these techniques is cost prohibitive for many potato breeding programs. X-ray data also lacks colorimetric features which may be needed to discriminate between different types of internal defect (bruising, greening, anthocyanin accumulation, internal brown spot) present within susceptible clones of breeding populations.

By far the greatest advantages of the technique presented is the low cost of deployment and potential to develop extensible models capable of classifying additional defects from the same data set. The capacity to inexpensively perform semiautomated, high resolution data collection, and downstream defect classification within potato breeding populations will enhance our ability to understand of how defect susceptibility is inherited and improve our ability to select productive, blemish free potato cultivars.

## VI. Conclusion

In this paper, a novel deep learning algorithm was proposed for potato hollow heart and non-hollow heart classification/detection. The performance of the proposed model was evaluated by investigating various metrics such as precision and the AUC. The results demonstrated the effectiveness of the model in different AWGN regions. As the next step, we aim to classify additional features (defects) from potato tubers. Many human scorable potato characteristics (sprouting, fungal/bacterial growth, bruising, and necrotic lesions) have the potential to be quantified using CNN classification algorithms. For future research, we aim to extend the proposed algorithms to detect and classify other potato defects such as internal brown spots, bruising, greening, and anthocyanin accumulation simultaneously. This requires constructing a potato data set that includes large numbers of each potato defect type. Therefore, additional data collection efforts are underway and we aim to determine if the proposed algorithms can be extended to classify these features.

## Acknowledgment

Computations supporting this project were performed on High-Performance Computing Systems at the University of South Dakota, funded by NSF Award 1626516. University of South Dakota (USD) Research Computing staff Addison Kleinsasser, Bill Conn, and Zach Hoiberg provided valuable technical expertise to this project.

