## Supplementary figures and images for "A deep learning algorithm for potato tuber hollow heart classification"

### Supplemental Fig. S1

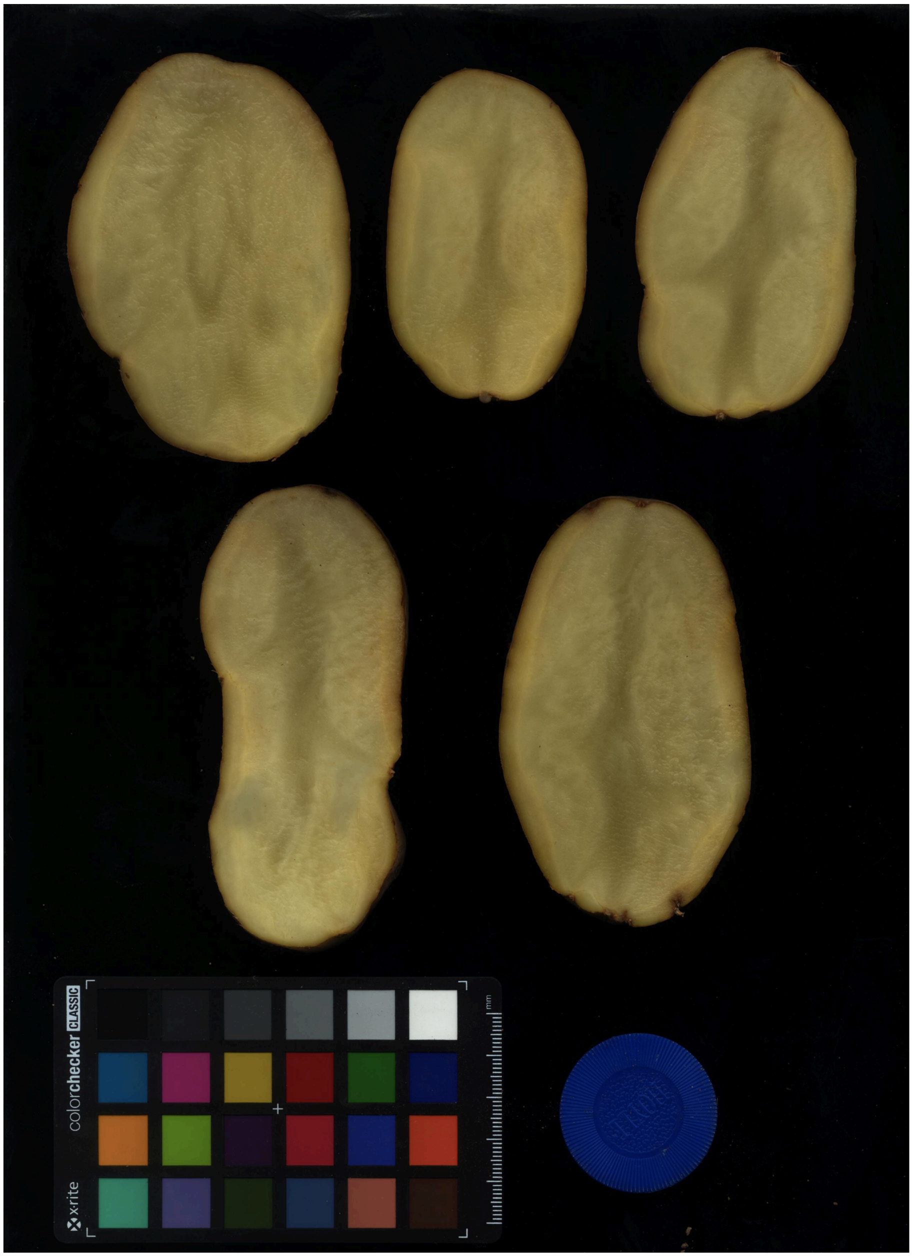


Fig. S1: Digital scans of tubers internal characteristics acquired using a flatbed scanner
